# Strong and lasting impacts of past global warming on baleen whale and prey abundance

**DOI:** 10.1101/497388

**Authors:** Andrea A. Cabrera, Elena Schall, Martine Bérubé, Lutz Bachmann, Simon Berrow, Peter B. Best, Phillip J. Clapham, Haydée A. Cunha, Luciano Dalla Rosa, Carolina Dias, Kenneth P. Findlay, Tore Haug, Mads Peter Heide-Jørgensen, Kit M. Kovacs, Scott Landry, Finn Larsen, Xênia Moreira Lopes, Christian Lydersen, David K. Mattila, Tom Oosting, Richard M. Pace, Chiara Papetti, Angeliki Paspati, Luis A. Pastene, Rui Prieto, Christian Ramp, Jooke Robbins, Conor Ryan, Richard Sears, Eduardo R. Secchi, Monica A. Silva, Gísli Víkingsson, Øystein Wiig, Nils Øien, Per J. Palsbøll

## Abstract

**Abstract:** The demography of baleen whales and their prey during the past 30 thousand years was assessed to understand the effects of past rapid global warming on marine ecosystems. Mitochondrial and genome-wide DNA sequence variation in eight baleen whale and seven prey species revealed strong, ocean-wide demographic changes that were correlated with changes in global temperatures and regional oceanographic conditions. In the Southern Ocean baleen whale and prey abundance increased exponentially and in apparent synchrony, whereas changes in abundance varied among species in the more heterogeneous North Atlantic Ocean. The estimated changes in whale abundance correlated with increases in the abundance of prey likely driven by reductions in sea-ice cover and an overall increase in primary production. However, the specific regional oceanographic environment, trophic interactions and species ecology also appeared to play an important role. Somewhat surprisingly the abundance of baleen whales and prey continued to increase for several thousand years after global temperatures stabilized. These findings warn of the potential for dramatic, long-term effects of current climate changes on the marine ecosystem.

*One Sentence Summary:* The effects of past global warming on marine ecosystems were drastic, system-wide and long-lasting.

## Main Text

The current global warming is affecting the distribution of species, population dynamics and trophic interactions across a wide range of ecosystems (*1*, *2*). However, predicting the long-term consequences of contemporary, short-term observations remains a challenge (*3*, *4*). The Pleistocene-Holocene transition was characterized by similarly rapid and dramatic increases in temperature and concomitant environmental changes (*4*, *5*) (Fig. 1), providing an opportunity to assess the long-term consequences of such changes in natural populations. Indeed, the effects of the Pleistocene-Holocene transition (7-12 thousand years ago, kya (*6*)) upon range shifts and extinction rates have been assessed in some detail in terrestrial megafauna (*7*, *8*), highlighting substantial spatio-temporal variation among and within species (*9*). In contrast, the effects of this period of dramatic climate change are poorly known for marine megafauna.

**Fig. 1.**
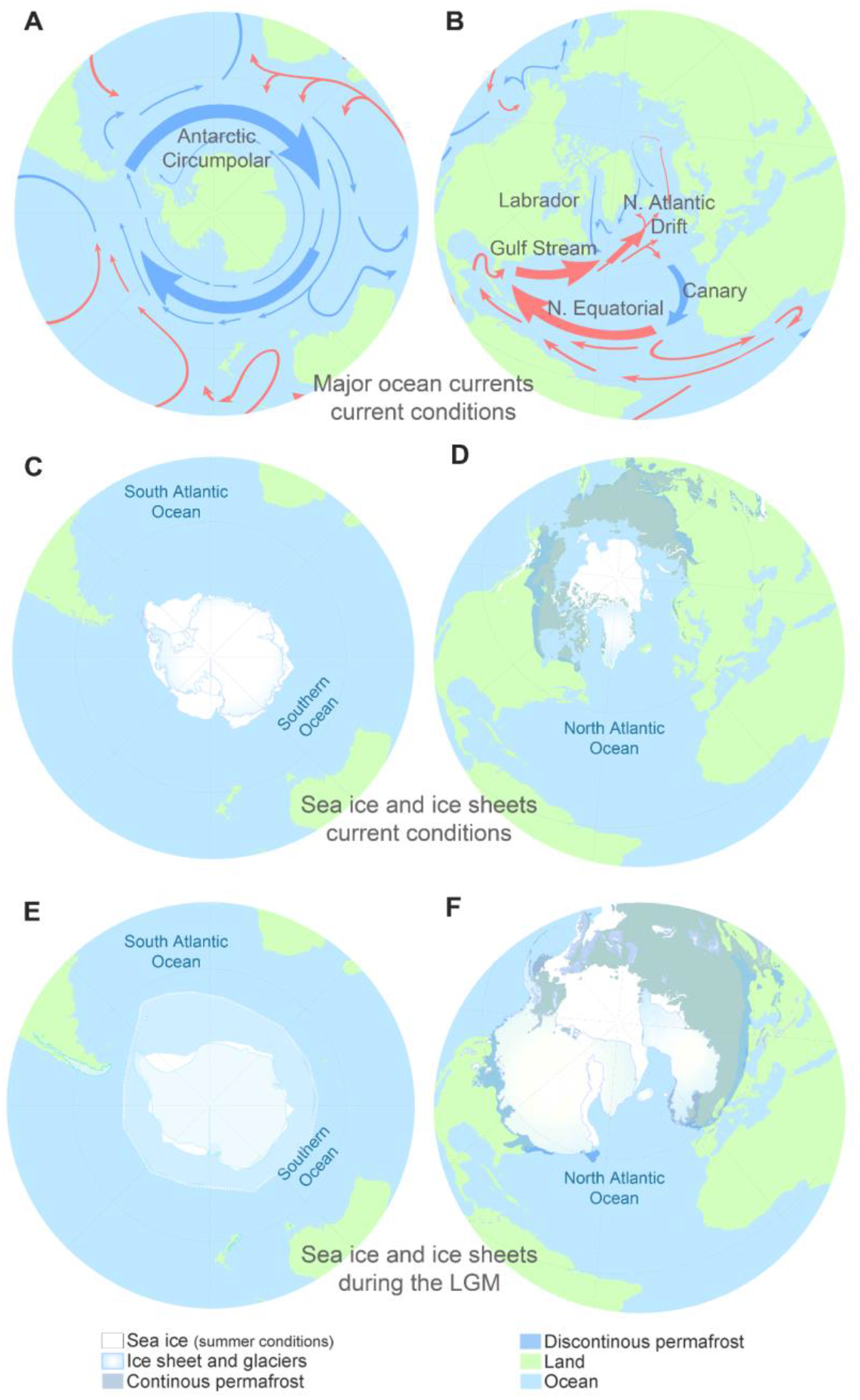
Major ocean currents and summer sea ice conditions before and after the Pleistocene-Holocene transition. (A - B) Simplified depiction of the major surface ocean currents in the Southern Ocean and North Atlantic Ocean. (C - D) Contemporary summer sea and land ice coverage. (E - F) Inferred summer sea and land ice coverage during the LGM.

Baleen whales are globally-distributed marine megafauna that feed on invertebrates and fish. Most baleen whale species undertake extensive seasonal migrations between low-latitude breeding grounds and high-latitude feeding areas (*10*). Consequently, baleen whales are subject to environmental and ecological changes across entire ocean basins. Here, DNA sequence data were employed to estimate demographic changes in eight baleen whale and seven prey species since the Last Glacial Maximum (LGM (19-26 kya (*5*)) until the late Holocene (~1 kya). The study spanned two ocean basins with contrasting large-scale oceanographic features (*11*) and included prey species in an attempt to elucidate the interplay between population dynamics, the environment and trophic interactions (*12*).

The Southern Ocean is a large ocean basin governed by the stable Antarctic Circumpolar Current (*13*) (Fig. 1a) and a pelagic food web dominated by Antarctic krill (*14*), resulting in a relatively homogenous marine ecosystem. In contrast, the North Atlantic is a much smaller ocean basin influenced by multiple, interacting ocean currents, continental run-off and cyclic climate oscillations (*15*, *16*) (Fig. 1b). The distribution and abundance of baleen whale prey in the North Atlantic Ocean varies markedly across time and space (*17*). The two oceans also differ by the relative reduction in sea ice cover since the LGM, which were comparatively more pronounced in the North Atlantic (Fig. 1c-f).

The coalescent-based Bayesian framework implemented in the software MIGRATE-N (*18*) was employed to infer temporal trends in genetic diversity (*θ*, the effective population size scaled by the generational mutation rate per nucleotide site, *μ*) and inter-ocean immigration rates (*M*, the immigration rate *m* scaled by *μ*) during the last 30 ky for each population. The parameters *θ* and *M* served as proxies for intra-ocean abundance and inter-ocean connectivity, respectively. The analyses targeted baleen whales as well as pelagic fish and invertebrate species that are preyed upon by baleen whales or at the same trophic level as baleen whale prey species (see Methods). In total, 4,761 mitochondrial DNA (mtDNA) sequences from eight baleen whale species and 2,271 mtDNA sequences from seven fish and invertebrate species were used for the estimations (Table S1). The data originated from specimens collected in the Southern Hemisphere (the South Atlantic Ocean, the southwestern Indian Ocean and the Southern Ocean) and in the North Atlantic Ocean (Fig. S1). Similar estimates of the temporal changes in *θ*, inferred from the folded site frequency spectrum were estimated from genome-wide SNP genotypes collected from 100 specimens in three baleen whale species, as a means to corroborate the estimates derived from the mtDNA sequences (Fig. S4, Supplementary information). In common with other studies based upon similar inference methods, all estimates of abundance and connectivity relied upon assumptions regarding mutation rates, which were subject to considerable uncertainty. Although the choice of mutation rates impacts the absolute values of estimates (i.e., abundance, connectivity and years) they do not impact the relative difference among estimates, which the inferences reported here were based upon.

The Pleistocene-Holocene transition was characterized by a steep rise in temperature, which, in turn, accelerated deglaciation rates, reduced sea ice cover and increased sea levels (*4*, *5*). The net result was an overall expansion of open water marine habitats, large-scale changes in ocean circulation and an overall increase in primary productivity (*4*-*6*) (Fig. 1). These changes were evident in the estimated increase in abundance across all baleen whale populations in both the North Atlantic Ocean and the Southern Hemisphere (Fig. 2). The finer-scale temporal trends in abundance were concomitant with the differences in oceanographic features between the North Atlantic and Southern Hemisphere ocean basins (*11*).

**Fig. 2.**
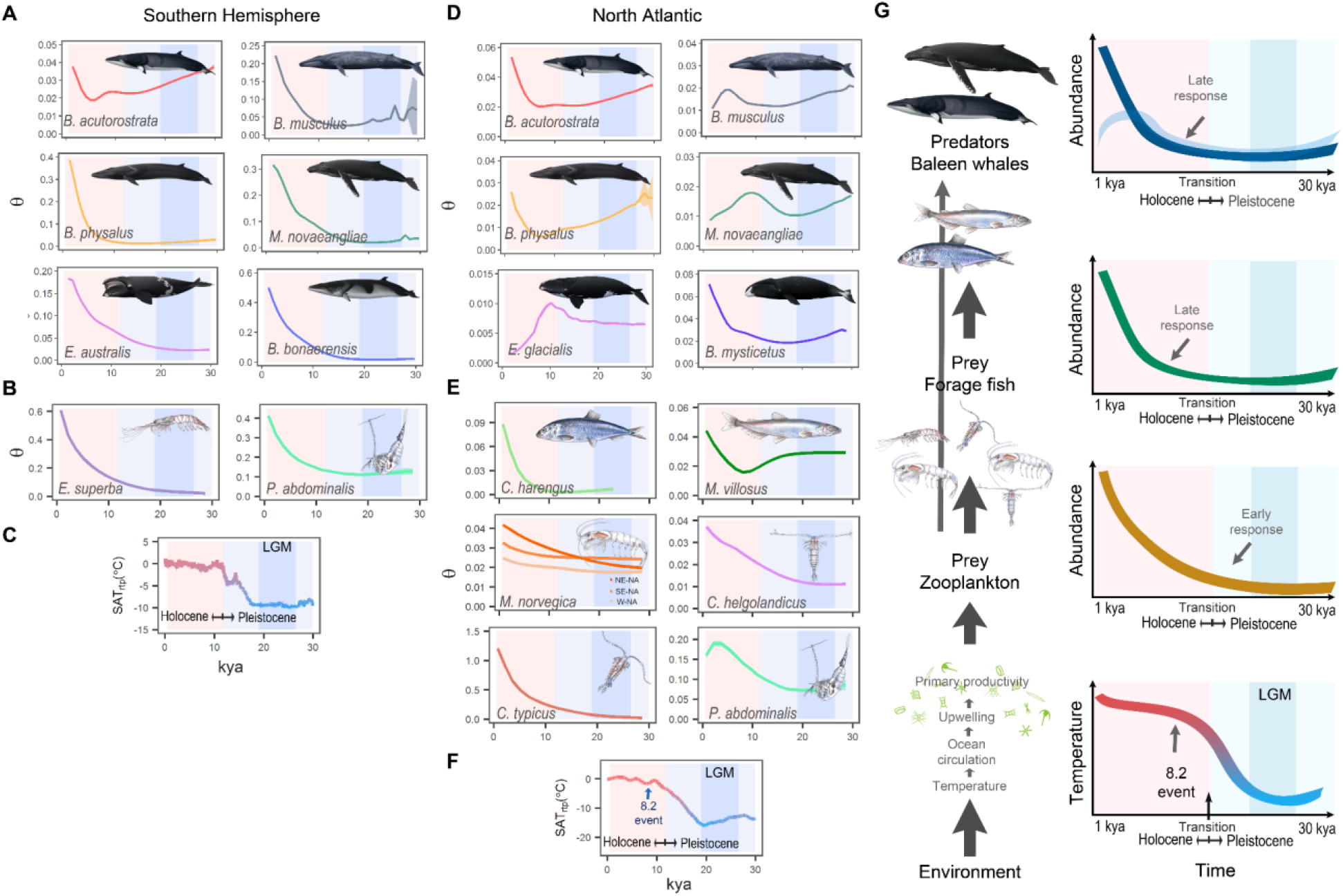
Estimated temporal trends of *θ* (as proxy for baleen whale and prey abundances) during the Pleistocene and Holocene (1 -30 kya). (A & D) Baleen whale species, (B & E) prey species. Note the different scales of the values on the vertical axis in genetic diversity (*θ*). Horizontal axis denotes the time in thousands of years ago (kya). (C & F) Historical surface air temperature relative to present temperature (SAT_RTP_) in degrees Celsius (°C). NE-NA: Northeastern North Atlantic (NA), SE-NA: Southeastern NA. W-NA: Western NA. (G) Graphic depiction of the bottom-up control of the demographic response of baleen whales during the Pleistocene-Holocene transition suggested by the results of this study. Red- and light blue-shaded areas indicate the LGM.

In the Southern Hemisphere, large, exponential and apparently synchronous increases in abundance were observed in all baleen whale species (Fig. 2a). These increases paralleled, similar large increases in prey abundance; i.e., Antarctic krill and copepods (Fig. 2b). The lagging increase in abundance of the common minke whale (*Balaenoptera acutorostrata*) likely reflects the difference in distribution and prey preferences of common minke whales. Common minke whales are mainly encountered at lower latitudes and feed on myctophid fishes, whereas the other baleen whale species feed mainly on krill at higher latitudes.

In contrast, North Atlantic baleen whale abundance trends were not synchronous to the extent observed in the Southern Hemisphere (Fig. 2d). The North Atlantic baleen whale species all underwent an initial expansion when temperatures increased, either immediately after the LGM (19-26 kya (*5*)) or later, during the initial phase of the Pleistocene-Holocene transition. However, the subsequent trends in abundance varied considerably among species. The blue whale (*B. musculus*), the humpback whale (*Megaptera novaeangliae*), and the North Atlantic right whale (*Eubalaena glacialis*) all underwent subsequent declines in abundance. The temporal trends in abundance also varied among the prey species. The abundance of northern krill (*Meganyctiphanes norvegica*) and copepods species (*Calanus helgolandicus*, *Centropages typicus*, *Pleuromamma abdominalis*) increased after the LGM, whereas the abundance of capelin (*Mallotus villosus*) and herring (*Clupea harengus*) did not increase until approximately 6-8 kya (Fig. 2e).

The relative change in baleen whale abundance (denoted Δ*θ*) at 1 kya relative to the 21 kya (i.e., at the LGM) was largest in the Southern Hemisphere, where the average Δ*θ* was estimated at 9.0 (range: 1.3-34, Fig. 3a). In comparison, the average Δ*θ* was estimated at 1.2 (range: 0.3-3.6, Fig. 3a) in the North Atlantic Ocean. A similar, albeit less pronounced, difference in average Δ*θ* between the two oceans was observed among prey species as well (Fig. 3b). This difference in Δ*θ* between the two oceans may, in part, stem from the proportionally larger increase in suitable baleen whale habitat and higher increase prey abundance in the Southern Ocean.

**Fig. 3.**
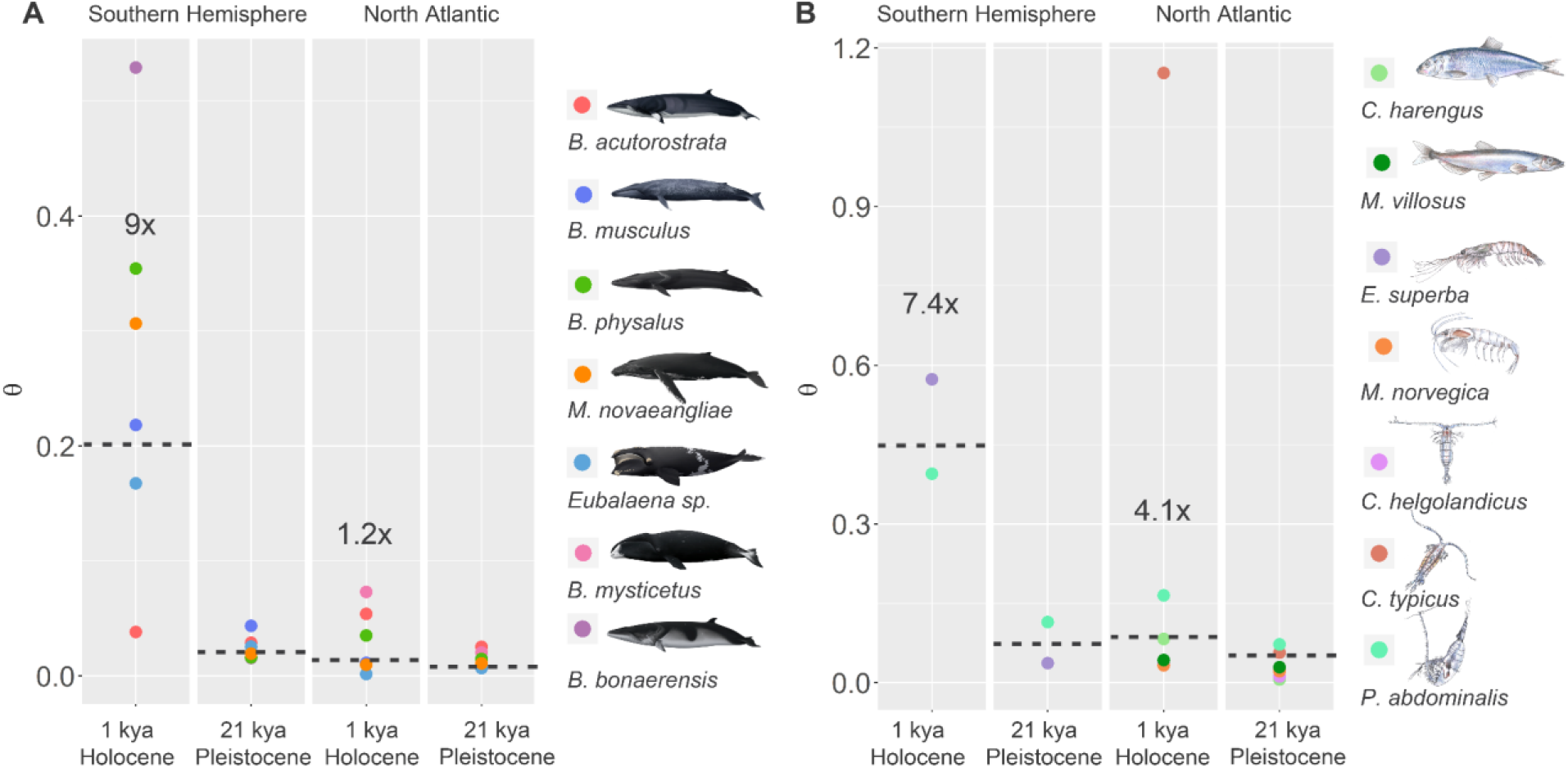
Estimated relative change in abundance for baleen whales and prey during the Pleistocene and Holocene. (A) Baleen whales and (B) prey species. Circles represent the median point estimates of *θ* in each species. Dotted lines indicate the geometric mean of *θ* (estimated from all point estimates). The numbers with an x (e.g., 7.5x) indicate the relative change in *θ* (Δθ) at one thousand years ago (kya) relative to 21 kya.

The estimates of inter-ocean basin connectivity were subject to high levels of uncertainty (Fig. S2). However, in general, baleen whale connectivity appeared to increase during two periods, when abundance peaked after the Pleistocene-Holocene transition and when suitable baleen whale habitat was greatly reduced and contracted towards the Equator during the LGM (Fig. 1c-f) bringing con-specific populations in close proximity.

The finding of similar increases in abundance at lower trophic levels (i.e., in key invertebrates, such as krill and copepods) suggests a bottom-up (*19*, *20*) enrichment of the oceans during the initial warming phase of the Pleistocene-Holocene transition (Fig. 2g). A parallel increase was particularly evident in the Southern Hemisphere where the trends in abundance among most baleen whales and Antarctic krill (*21*) were strongly correlated (r=0.88-1.00, p<0.005, Fig. S3). This result was consistent with previous paleo-oceanographic models that suggested an increase in primary productivity during the Pleistocene-Holocene transition (*22*, *23*), which was characterized by a shift in phytoplankton composition from perennial pelagic to seasonal sea-ice-associated species. The latter species are viewed as indicative of high levels of primary productivity (*24*, *25*).

The trends in abundance in the North Atlantic Ocean during the Pleistocene-Holocene transition varied across space and time. The inferred abundance trajectories of most baleen whales, fish and some copepod species in the North Atlantic Ocean changed markedly around 6-8 kya (Fig. 2). A possible cause of such ocean-wide changes was the 8.2 kya event, when global ocean temperatures dropped precipitously due to a massive discharge of glacial melt water into the western North Atlantic Ocean from proglacial lakes (*26*, *27*). This event led to a shift in phytoplankton composition consistent with a reduction in primary productivity, in particular in the western North Atlantic (*24*, *25*).

Recent predictions of the effects of the current global warming on marine mammal populations have relied upon field observations (*28*, *29*). Baleen whale species, such as humpback, fin (*B. physalus*) and blue whales, appear to arrive earlier and at higher latitudes on the summer feeding grounds, increasing competition with polar species, such as the bowhead whale (*Balaena mysticetus*) (*30*). However, although some baleen whale species appear to benefit from global warming at present, the findings reported here suggest that the oceanographic and ecological changes introduced by global warming initiated geological and biological processes with long-lasting and wide-ranging impacts on the marine ecosystem. Even though the rapid rise in global temperatures during the Pleistocene-Holocene transition plateaued ~10 kya, most vertebrate and invertebrate species targeted in this study continued to increase in abundance in both hemispheres until 1 kya, the most recent time included in the analysis. In other words, the Pleistocene-Holocene transition set in motion long-term oceanographic and ecological changes that continued to affect both abundance and connectivity of baleen whales and their prey for ~10 ky. Consequently, current global warming is likely to exert similarly drastic, long-term and wide-ranging changes on marine ecosystems even after temperatures have stabilized. Accordingly, projections of impacts of global warming on marine species should account for both the short and long-term effects as well as the complexity of the oceanographic and ecological interactions evident in this study.

## Supporting information

## Acknowledgments

The baleen whale illustrations were reproduced with the permission of Frédérique Lucas and prey illustrations were drawn by Ligia Arreola. Hans J. Skaug kindly provided sample material. Steve Beissinger, Robin Waples and Richard Hudson provided helpful suggestions.

## Author contributions

PJP and AAC conceived and designed the study with input from ES. LB, SB, PBB, PJC, HAC, LDR, KPF, TH, MPH, KMK, SL, FL, CL, DKM, RMP, CP, LAP, CR, JR, CR, RS, ERS, MAS, RP, HJS, GV, ØW, NØ provided data or sample material. MB, TO, CD conducted laboratory analyses. AAC, ES, XML conducted data analyses supervised by PJP and with input from AP. AAC designed figures. AAC and PJP drafted the manuscript with contributions from MB, ES, JR, CR, PJC and XML. All authors read, edited, commented on and approved the final manuscript;

## Supplementary Materials

Materials and Methods

Figures S1-S7

Tables S1-S5

References

