## Supplementary material for "Strong and lasting impacts of past global warming on baleen whale and prey abundance"

**This PDF file includes:**

Materials and Methods
Figs. S1 to S7
Tables S1 to S5
References

### Materials and Methods

Taxon selection

The study focused on eight species of baleen whale as well as seven fish and invertebrate prey species (Table S1). The common minke whale (*B. acutorostrata*), the blue whale (*B. musculus*), the fin whale (*B. physalus*), and the humpback whale (*M. novaeangliae*) all have global distributions (Fig. S1). The North Atlantic right whale (*E. glacialis*) and the southern right whale (*E. australis*), are limited to the North Atlantic and Southern Hemisphere, respectively. In the analysis, these two nominal species were treated as different ocean basin populations of a single species in light of their low degree of genetic divergence (*31*) ($F_{ST}$: 0.21 (*32*)) which is similar to the inter-ocean differences among other baleen whale populations (e.g., North Atlantic, North Pacific and Southern Hemisphere humpback whales ($F_{ST}$: 0.151-0.516)) (*33*). The remaining two baleen whale species have restricted distributions (Fig. S1b); the bowhead whale (*B. mysticetus*) is confined to the Arctic, and the Antarctic minke whale (*B. bonaerensis*) to the Southern Ocean, both species, with a few recent, notable exceptions (*34-36*).

The fish and invertebrate species included two krill, three copepod and two small schooling fishes. These represent known baleen whale prey (i.e., Antarctic krill, (*E. superba*), northern krill (*M. norvegica*), the copepod (*C. typicus*), North Atlantic herring (*C. harengus*) and capelin (*M. villosus*)) or occupy the same trophic level as similar baleen whale prey (i.e., two copepod species (*C. helgolandicus* and *P. abdominalis)* in place of *Calanus finmarchicus*, which is the prey of right and bowhead whales; Table S1). Antarctic krill is confined to the Southern Ocean. The northern krill, two copepod species (*C. typicus*, *C. helgolandicus*), the North Atlantic herring and the capelin are only found in the Northern Hemisphere. The copepod (*P. abdominalis*) has a global distribution (Fig. S1).

Sample collection

In case of novel DNA sequence data generated specifically for this study, skin samples from baleen whales (Table S1) were collected from free-ranging individuals using remote biopsy sampling approaches (*37*) or from dead individuals.  The latter included naturally stranded whales bycatch and whaling operations conducted prior to the current international moratorium or as part of aboriginal subsistence whaling (permitted under the agreements of the International Whaling Commission). All tissue samples were stored in a solution of saturated sodium chloride and 20% dimethyl sulfoxide and archived at -20 or -80 degrees Celsius (°C) until analysis. Total-cell DNA was extracted from the tissue samples using either standard phenol and chloroform extraction (*38, 39*) or DNeasy™ columns (Qiagen Inc.) following the manufacturer's instructions.

Multi-locus microsatellite genotypes (data not included) were employed to identify and remove duplicate samples of the same individual as well as mother and calf pairs sampled during the same sighting event (i.e., non-independent samples of closely related individuals). Closely related individuals sampled at random (i.e., during different sighting events) were not excluded from the analysis (*40*).

Mitochondrial DNA data

In total, 4,761 baleen whale mitochondrial control region DNA sequences and 2,271 DNA sequences from fish and invertebrate species were included in the study. The latter DNA sequences were from different mtDNA genes, such as; cytochrome *c* oxidase subunit I (COI); NADH dehydrogenase subunit 1 (ND1); cytochrome *b* (CYTB); or 16S ribosomal DNA (16S rDNA; Table S1). The mtDNA sequence data were either generated during this study or obtained from published sources (*33, 41-54*). The experimental conditions for published data were described in the relevant publications (Table S1). New mtDNA sequence data were generated as described below.

*Laboratory methods*: The first ~400 base pairs (bp) of the 5' end of the mitochondrial control region were amplified, using the DNA oligo-nucleotides MT4F(*55*) and BP16071R(*56*). The initial PCR amplifications were performed in a 20 µL reaction volume consisting of 0.2 µM of each dNTP, 67 mM Tris-HCl (pH 8.8), 2 mM $MgCl_2$, 17 mM $NH_3SO_4$, 10 mM $\beta$-mercaptoethanol, 0.1 µM of each DNA oligo-nucleotide, 0.4 units of *Taq* DNA polymerase (Fermentas Inc.) and ~10 - 20 ng of extracted DNA. The thermo-cycling conditions were: 2' (minutes) at 94 °C, followed by 25 cycles each with of 15'' (seconds) at 94 °C, 30'' at 54 °C and 120'' at 72 °C. Unincorporated nucleotides and excess primers were removed using shrimp alkaline phosphatase and exonuclease I (*57*). Cycle sequencing was conducted according to the manufacturer's instructions (1/16$^{th}$ the recommended amount of Big Dye™ v3.1 Terminator Ready Reaction Mix, Life Technologies Inc.) with the DNA oligo-nucleotides MT4F or BP16071R. Excess nucleotides were removed by ethanol precipitation(*58*) and the cycle-sequencing products re-suspended in 10 µL deionized formamide (Calbiochem Inc.). The order of cycle-sequencing products was resolved by capillary electrophoresis (Applied Biosystems ABI Prism™ 3730, Life Technologies Inc.). The quality of DNA sequence chromatograms were inspected by eye using the software CHROMAS® (ver. 2.13, Technelysium Pty Ltd.) or SEQUENCHER® (ver. 5.1, Gene Codes Corporation).

*Data processing and sequence alignment:* DNA sequence alignment was performed using the ClustalW algorithm (*59*) and default parameter settings as implemented in MEGA (ver. 6.0)(*60*) followed by visual inspection of the alignment. All sequences were trimmed to equal length (Table S1).

*Estimation of temporal trends of genetic diversity and migration rates from single-locus DNA sequences:* The software MIGRATE-N (ver. 3.6.6) (*61, 62*) was employed to estimate skyline plots of temporal changes in regional genetic diversity ($\theta$) and immigration rates scaled by generational mutation rate per nucleotide site (*M*). The software JMODELTEST (ver. 2) (*63*) was employed to select the most probable nucleotide mutation model and mutation parameter values (Table S2). The prior ranges of $\theta$ and *M* were determined from preliminary estimations with reduced sample sizes and short Monte Carlo Markov chains (MCMC) using the estimates of the $F_{ST}$ as starting values. The priors were then adjusted according to the outcomes of the preliminary estimations. The specific analysis parameter values employed during the final estimations are listed in Table S2. For data sets with more than 250 DNA sequences, random sub-sampling (without replacement) was employed at sample sizes of 250 DNA sequences per sample partition. In the cases of data sets that differed in regional sample sizes by more than 100% (i.e., common minke and fin whales), a random sub-sampling equal to the size of the smallest sample partition was employed, i.e., 23 and 61 for the common minke whale and the fin whale, respectively (see notes on "effect of sample size", Fig. S6). Final estimates were based upon three independent MCMC estimations, all initiated with different random seeds. Each MCMC run comprised 100 replicates, each consisting of a single long MCMC of eight million burn-in steps followed by an additional eight million steps, sampled at every 200$^{th}$ step. A static heating scheme of four chains at temperatures 1.0; 1.5; 3.0 and 1,000,000, respectively, was employed. Convergence was assessed employing the R-CRAN package CODA (*64*). Consistency among the three MCMC estimations, smooth and unimodal distribution within the prior range for all estimates were also considered as indications of convergence. The effective sample sizes of all MCMC estimations were larger than 10,000. The final point estimates and standard deviation (~95% CI) of $\theta$ and $M$ per time interval were obtained by combining the results of the three independent MCMC estimations. In order to balance the different number of MCMC data points employed to infer $\theta$ and $M$ in each of the three independent MCMC estimations, a weighted average of the medians and standard deviations of $\theta$ and $M$ was estimated (i.e., pooled median and pooled standard deviation) per time interval. The pooled median of $\theta$ and $M$ was estimated as: $m_p = \sum_{i=1}^{k} n_i\, m_i / \sum_{i=1}^{k} n_i$, where, $n_i$ denotes the number of MCMC data points employed in estimate $i$, $m_i$ denotes the estimated median parameter value of estimate $i$, $i$ denotes the estimated parameter value of $\theta$ or $M$, $k$ denotes the total number of estimated parameter values (i.e., three independent MCMC estimations). The pooled standard deviation was estimated as $SD_p = \sqrt{\sum_{i=1}^{k}(n_i - 1)SD_i{}^2 / ((\sum_{i=1}^{k} n_i) - k)}$, where $SD_i$ denotes the estimated standard deviation of estimate $i$. The 95% credible interval of the final estimations was approximated to $m_p \pm 1.96\, SD_p/\sqrt{n_i}$ assuming $m$ was normally distributed.

Possible effects of spatial population genetic structuring within the North Atlantic Ocean or the Southern Hemisphere were assessed by comparing the outcome of analyses based upon pooled and spatially separate samples. Population samples were analyzed separately in those cases where a discernible effect of spatial population genetic structure on the final estimates was detected (e.g., *M. norvegica*).

The conversion of time estimates (which were scaled against $\mu$) to years required estimates of generational mutation rates. A range of reported mutation rates in the targeted or closely related species was explored (see "notes on mutation rate"). Different generational mutation rates in the non-coding (i.e., control region) and the coding genes (i.e., COI, ND1, CYTB and 16S rDNA) in the mitochondrial genome were considered, but similar generational mutation rates among species were assumed for the same gene. In the case of non-coding genes, a generational mutation rate at 1.125 x $10^{-6}$ per bp was applied, which was within the range of previously estimated values (from 2 x $10^{-7}$ to 2 x $10^{-5}$ bp, Fig. S7). The generational mutation rate was based upon the annual mutation rate at 5.3 x $10^{-8}$ per bp reported by Alter and Palumbi (*65*) for the common minke whale and a generation time at 21.2 years, (the average of values estimated by Pacifici *et al*. (*66*) and Taylor *et al*. (*67*) for the common minke whale). In the case of coding mitochondrial genes, a generational mutation rate at 3.4 x $10^{-7}$ per bp was applied, also within the range of reported values of generational mutation rates from mitochondrial coding regions or for entire mitochondrial genomes (from 2 x $10^{-8}$ to 2 x $10^{-4}$ per bp, Fig. S7). This value was derived using from an estimate of the annual mutation rate at 1.7 x $10^{-8}$ for the coding parts in the human mitochondrial genome (*68*) and a generation time of 20 years (*68, 69*). This generational mutation rate was similar to other, direct estimates of the generational mutation rate for the entire mitochondrial genome reported in several invertebrate model species (*70, 71*).

The estimation period was limited to one kya to 30 kya, in order to include the LGM and exclude possible recent anthropogenic effects, such as whaling.

The consistency of the results obtained from different mitochondrial genes was assessed by comparison of the temporal trends of $\theta$ to estimates of different mitochondrial genes from the same species (Tables S2-S3, Fig. S4). Con-specific temporal trends in abundance estimated from mitochondrial DNA sequences were also compared to the estimates based on genome-wide SNPs (single nucleotide polymorphisms) generated by next generation sequencing in three selected baleen whale species as described below (see Nuclear DNA data, Table S3, Fig. S4).

Mutation rates

*Choice of mutation rate.* The estimates of time ($\tau$) obtained from the coalescent-based approach employed in this study are scaled by the generational mutation rate per nucleotide site ($\mu$). Consequently, a conversion of $\tau$ into years ($t$) was necessary in order to relate the estimated temporal trends in $\theta$ to environmental change, such as increasing global temperatures. Such conversion required an estimate of the generation time or an annual mutation rate per site for the DNA sequences from which $\tau$ was estimated. The estimates of temporal trends in abundance, or more precisely the effective population size ($N_e$), were inferred from $\theta$ ($= 4N_e\mu$) as the focus was on relative (as opposed to absolute) change in $N_e$. Relying upon $\theta$ assumes that $\mu$ is constant, an assumption that some authors have argued may not necessarily hold true (*72, 73*).

Mutation rates vary considerably among different DNA sequences across a genome(*74*), as well as among genomes and species (*75*). Additionally, reported mutation rates also vary among estimation methods, by as much as an order of magnitude (e.g., *76, 77, 78*). Examples of published mutation rate estimates based upon coding and non-coding mtDNA sequences from species targeted in this study (or from the same taxonomic group when the targeted species were not available) are listed in Table S4 and S5.

In this study, different generational mutation rates were applied for coding and non-coding DNA sequences (*79*) whereas the same ***generational*** mutation rate was applied across all species for coding and non-coding DNA sequences, respectively. In terms of the ***annual*** mutation rate, this choice implied that species with short and long generation times have high and low annual mutation rates, respectively. The consistency of the choice of the generational mutation rates for non-coding and coding DNA sequences was corroborated by a comparison of estimates of the trends of $\theta$ obtained from different DNA sequences in the same species (Fig. S7c), although such an assessment did not negate the possibility of a systematic bias.

*General logic in choices of mtDNA mutation rates.* The estimates in this study were (in part) based upon sequence variation in non-coding and coding mtDNA sequences. The specific choice of DNA sequences was determined by data availability (published or new data). All mtDNA data from the baleen whales were from the non-coding control region. The mtDNA sequence data from fish and invertebrate species were from published sources and all from coding mtDNA sequences, such as; cytochrome *c* oxidase subunit I (COI); NADH dehydrogenase subunit 1 (ND1); cytochrome *b* (CYTB); or 16S ribosomal DNA (16S rDNA).

As mentioned above, different generational mutation rates were applied for coding and non-coding mtDNA(*79*). The specific mutation rates were chosen from species with well-characterized generation times and available estimates of annual mutation rates; or species from which estimates of the generational mutation rate were available. Among marine fishes and invertebrates, it has been commonplace to employ annual mutation rates that used the rise of Central American Isthmus as the temporal calibration point (*80*). However, recent findings have suggested that the conventionally recognized timing of the closure of the Central American Isthmus is possibly incorrect (*81, 82*), and hence mtDNA mutation rate estimates based solely upon this calibration point were excluded. In the case of the coding mtDNA sequences, the applied mutation rate was selected from those species where an average mutation rate for multiple coding mtDNA sequences was available, i.e., excluding estimates based upon a single coding mtDNA sequence or the entire mitochondrial genome (i.e., including the faster evolving non-coding DNA sequences).

*Specific mutation rates.* Following the rationale outlined in the above sections, specific mutation rates were chosen as described below.

One generational mutation rate of $1.12x10^{-6}$ per base pair (bp) was applied to all baleen whale estimations based upon mtDNA control region sequences. The applied generational mutation rate for the baleen whale mtDNA control region was obtained from the annual mutation rate estimated from minke whale (*Balaenoptera acutorostrata*) mtDNA control region sequences (i.e., 5.3 x $10^{-8}$ per bp) (*65*) and a generation time of 21.2 years (*66*, *67*). This rate fell within the range of previously applied values (from 2 x $10^{-7}$ to 2 x $10^{-5}$ per bp, median value: 1.24 x $10^{-6}$ per bp, Table S5, Fig. S7a).

Similarly, one generational mutation rate (3.4 x $10^{-7}$ per bp) for coding mtDNA sequences was applied to all invertebrate and vertebrate species. The specific rate applied was based upon an estimated annual mutation rate at 1.7 x $10^{-8}$ for the coding mtDNA sequences in the human mitochondrial genome reported by Ingman *et al*.(*68*) and a human generation time of 20 years(*69*). This rate fell within previously reported values from vertebrate and invertebrate species (from 2.5 x $10^{-8}$ to 1.2 x $10^{-6}$ (with one exception at 2.0 x $10^{-4}$ per bp), median value: 1.03 x $10^{-7}$, Table S5, Fig. S7b).

The consistency of the estimated trends and time points of change in $\theta$ between non-coding and coding mtDNA sequences was assessed by comparing estimates from non-coding and coding mtDNA in North Atlantic herring (*Clupea harengus*) for which such data were available (Table S2-S3). As expected, the estimates were consistent among non-coding and coding mtDNA using the applied annual mutation rates.

Effect of sample size

The results obtained from the mtDNA sequences of six baleen whales were compared to two values of sample size per partition (i.e., 250 and 23, Fig. S6). In the first case of a random sub-sampling at 250 mitochondrial DNA sequences per partition, three species had uneven sample sizes (i.e., one of the sampled populations had less than 250 mitochondrial DNA sequences) and two species had even sample sizes (Fig. S6a). For the three species with uneven sample sizes, two differed by more than 100% (i.e., common minke whale and fin whale). In the second case, a random sub-sampling equal to the size of the smaller sample partition from all species (i.e., 23) was employed. The sample size of the smaller sample partition was selected in order to compare all species with equal and even sample sizes.

In general, the temporal trends in $\theta$ based on a random sub-sampling at 250 mitochondrial DNA sequences were consistent with those estimated from the mtDNA sequence variation of a random sub-sampling at 23 mitochondrial DNA sequences (Fig. S6). The consistency between large and small sample sizes in coalescent-based estimations was previously noted by Plutzhnikov and Donnelly (*83*) and Felsenstein (*84*), who found that a small number of samples (8) is sufficient to accurately inference of $\theta$ in a single isolated population of fixed size. However, larger sample sizes are likely required in the presence of migration or population size changes (*83*).

Low levels of variation was observed in the case of the common minke whale. The median estimate of the temporal trend in $\theta$ based on uneven sampling indicated a lower level of recent population expansion (Fig. S6a) than was the case for the median estimate of the temporal trend in $\theta$ based on even sample sizes (Fig. S6b). In a coalescent-based approach, the estimate of a parameter relies on the completeness of the sampling across the entire parameter space during the MCMC estimation. Uneven sample sizes may result in poor sampling from populations with low sample sizes if the MCMC runs are not long enough (*85*). The diminishing gain in information at large sample sizes at the cost of poorly evaluated parameter estimates under uneven sampling conditions favored a sampling strategy with even population samples generated by random sub-sampling from large data sets (Methods, Fig 2).

Nuclear DNA data

*Laboratory methods:* Genome-wide SNP genotypes were generated from double digested restriction-associated (ddRAD) (*86*) and quaddRAD libraries (*87*). Common minke whale and Southern right whale libraries were generated from ddRAD libraries prepared as described by Peterson *et al. (86).* Fin whale libraries were generated from quaddRAD libraries prepared as described by Franchini *et al. (87).* All libraries were prepared from genomic DNA digested with *HindIII and MspI* and insert sizes between 300 and 400 bp. Libraries were sequenced on an Illumina HiSeq™ 2500 platform (ver. 4) at 100 (ddRAD) and 125 (quaddRAD) bp, paired-end sequencing with 10% PhiX spike-in.

*Data processing:* In the case of the quaddRAD library, PCR clones were removed using the *clone_filter* script implemented in the software suite STACKS (ver. 1.47) (*88*). Illumina HiSeq sequence data from both, quaddRAD and ddRAD, were de-multiplexed with *process_radtags* employing STACKS (ver. 1.47) with default settings. The filtered reads were aligned to a reference genome using BOWTIE2 (ver. 2.2.8) (*89*) as "end to end" alignment employing the setting *very_sensitive* (i.e., *D* 20, *R* 3, *N* 0, *L* 20 and *i S*,1,0.50). The maximum fragment length for paired-end alignments was 600 bp and discordant alignments were prohibited. In the case of the common minke whale and fin whale data, the common minke whale genome (*90*) was employed as a reference. The bowhead whale genome (*91*) was used as the reference in the case of the southern right whale.

The folded site frequency spectrum (SFS) (*92*) was estimated using ANGSD (*93*) from samples with a minimum of three million reads. SNPs with a base quality score below 20 and a mapping quality below ten were discarded. Only SNPs genotyped in a minimum of 80% of the individuals were retained in the final estimation. Two minimum coverage values (x10 and x2) were evaluated (see "notes on genome wide SNP genotype analyses"). SNP genotype frequencies were estimated using likelihood procedure implemented in GATK (*94*).

*Estimation of temporal trends of genetic diversity from genome-wide SNP genotypes:* The method implemented in the software STAIRWAY PLOT (ver. 2.0 beta) (*95*) was employed to infer the temporal changes in $\theta$ from the folded SFS estimated from the genome-wide SNP data. Two-thirds of the data were employed as training data and $(n_{seq}) - 2/4$, $(n_{seq}) - 2/2$, $3\ (n_{seq} - 2)/4$, and $n_{seq} - 2$ , where $n_{seq}$ was two times the sample size, as the number of random breakpoints. The optimal number of random breakpoints was selected based upon the results from the training data. In order to convert the time-scaled mutation rate estimates in years, an annual mutation rate of 1.07 x $10^{-9}$ per bp (*90*) estimated for the whole-genome of baleen whales was applied to all three baleen whale species. The average generation times estimated by Taylor *et al.* (*67*) and Pacifici *et al.* (*66*) were applied, i.e., 21.2 years for the common minke whale, 32.5 years for the fin whale and 27.6 years for the southern right whale. The estimations were limited to the period of 1-30 kya.

Notes on genome-wide SNP genotype analyses

The results obtained from the mtDNA sequences in three baleen whales were corroborated with similar estimates of the temporal trends in $\theta$ based upon genome-wide SNP genotypes in a total of

100 baleen whale specimens across three species. The temporal trends in $\theta$ based on the genome-wide SNP genotypes were consistent with those estimated from mtDNA sequence variation (Fig. S4). The temporal trends of $\theta$ estimated from mtDNA sequences and the x2 coverage genome-wide SNP genotypes all suggested population expansions during the Pleistocene-Holocene transition in the three baleen whale species (i.e., the North Atlantic common minke whale (*B. acutorostrata*), the fin whale (*B. physalus*), and the southern right whale (*Eubalaena australis*)).

Increasing the read depth from x2 to x10 resulted in a less dramatic expansion. However, the temporal trend in the changes of $\theta$ remained the same for the common minke whale and the southern right whale at both levels of coverage. However, in the case of the fin whale, the median estimates of the temporal trend in $\theta$ at a coverage at x10 did not indicate a population expansion (Fig. S5). Although the median estimate did not indicate a population expansion, the confidence band included the possibility of an expansion and was very similar to the band inferred at a coverage at x2. Detection of recent expansions relies upon the presence of rare alleles, which in turn requires a large number of SNP genotypes and large sample sizes. A recent simulation-based evaluation demonstrated the inability of such "stairway plots" of temporal changes in $\theta$ to detect recent population expansions (i.e., <10 kya) when sample sizes were small (i.e., ~ 30 samples) (*95*). The sample sizes employed in this study were 27, 45 and 28 samples for the common minke whale, southern right whale and fin whale, respectively.

<u>Temperature data</u>

Surface air temperature (SAT) estimates for the Southern Hemisphere were inferred from deuterium measurements collected from the Antarctic EPICA Dome C Ice Core (*96*). For the Northern Hemisphere, continental atmospheric temperatures between 40 and 80° N calibrated with oxygen isotope records from 57 sediment cores were obtained from Bintanja *et al.* (*97*).

<u>Maps of ocean circulation and sea ice reconstructions</u>

Maps were generated with ArcGIS® (ver. 10.3, ESRI® Inc.). Ocean current data were obtained from the NOAA National Weather Service (*98*). Contemporary and LGM permafrost and ground ice data were obtained from Brown *et al.* (*99*) and Lindgren *et al.* (*100*), respectively. Average sea ice coverage during March and September 2016 (obtained from the National Snow and Ice Data Center (*101*)) represented contemporary summer sea ice coverage in the Antarctic and the Arctic, respectively. Contemporary ice sheet and glacial projections were obtained from Natural Earth (*102*). The data for the Antarctic summer sea ice coverage, ice sheet cover and extensions of the glaciers during the LGM were obtained from Gersonde *et al*., (*103*) and CLIMAP 1981 (*104*). The data for the Arctic summer sea ice coverage, ice sheet and extension of the glaciers from the LGM were obtained from GLAMAP 2000 (*105*) and Ehlers et al., (*106*). Maps were displayed using a South Pole or North Pole Lambert Azimuthal Equal Area projection and a World Geodetic System 1984 with map datum at a scale of 1:65,000,000.

<u>Maps of approximate species ranges</u>

Maps were generated with ARCGIS® (ver. 10.3, ESRI® Inc.). The approximate current species ranges for baleen whales, Antarctic krill and herring were obtained from the IUCN Red List of Threatened Species™ (*107*). The approximate current species ranges of capelin and northern krill were generated based upon data from AquaMaps (*108*). In the case of the three copepod species, approximate ranges were generated based on Bonnet *et al*. (*109*) and COPEPEDIA (*110*). Maps were displayed using a Miller cylindrical projection and a World Geodetic System 1984 with map datum at a scale of 1:15,000,000.

Correlation among baleen whales, prey and climate

Pearson's correlation coefficients were estimated using R (ver. 3.2.5) (*111*). Estimates of $\theta$ and SAT were fitted to 1,000 year intervals by linear interpolation as implemented in R CRAN (ver. 3.2.5) (*111*). Time intervals with missing data were excluded from the analysis. Consistency, and possible dependence issues, of the estimated correlations were evaluated employing different time intervals (i.e., every 1,000; 2,000 and 5,000 years) between data points (Fig. S3).

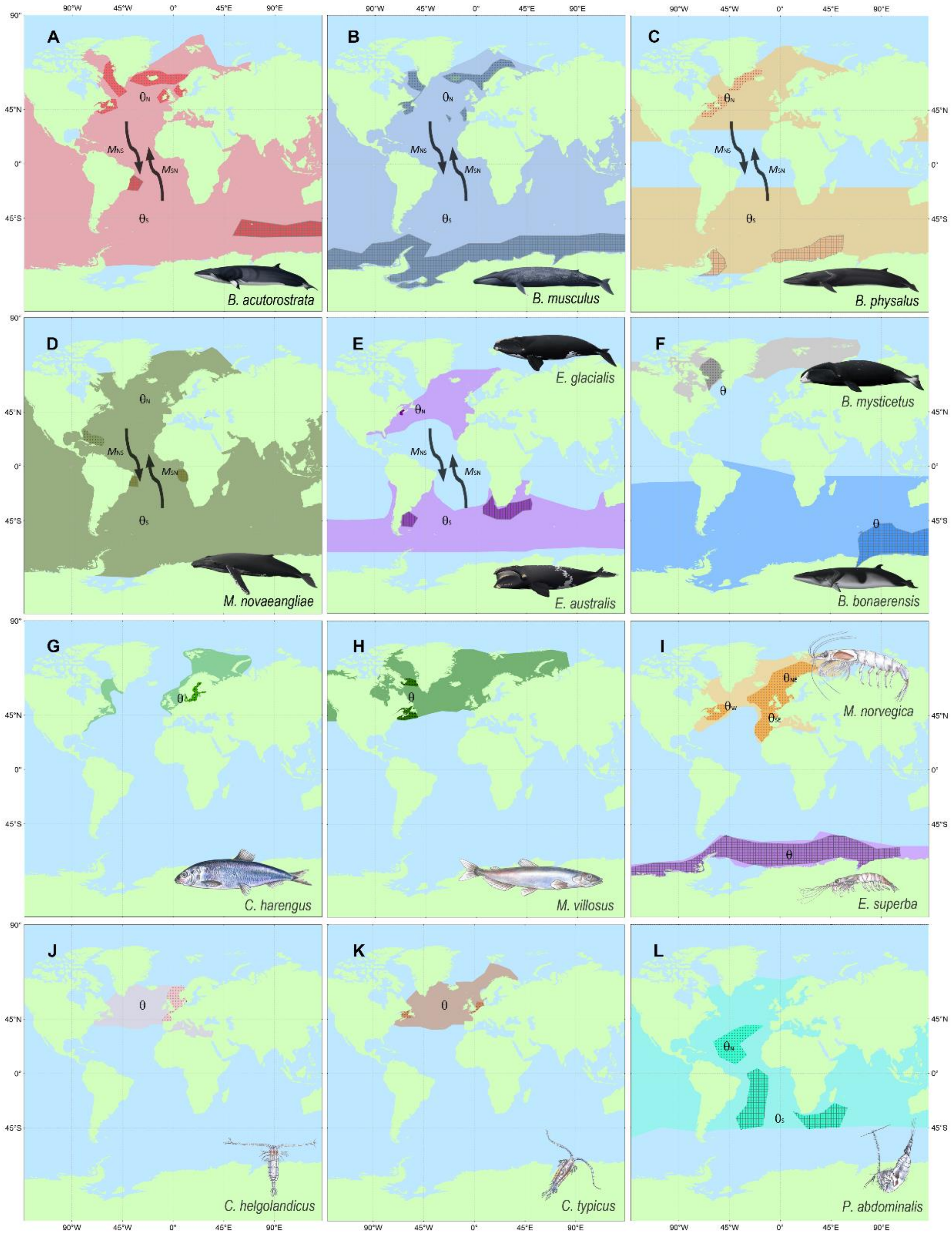


**Fig. S1.**

**Sampling locations and estimated parameters for North Atlantic and Southern Hemisphere baleen whales and prey species.** Current approximate species range and sampling location for (A - F) baleen whales: (A) common minke whale (B) blue whale, (C) fin whale, (D) humpback whale (E) North Atlantic and Southern right whale, (F) bowhead whale (top), Antarctic minke whale (bottom) and (G-L) prey species: (G) herring, (H) capelin, (I) northern krill (top) Antarctic krill (bottom), (J - L) copepod species. Estimated parameters $\theta$ ($\theta_N$: from the North Atlantic populations and $\theta_S$: from the Southern Hemisphere populations) and $M$ ($M_{NS}$: from the North Atlantic Ocean into the Southern Hemisphere, $M_{SN}$: from Southern Hemisphere into the North Atlantic Ocean).

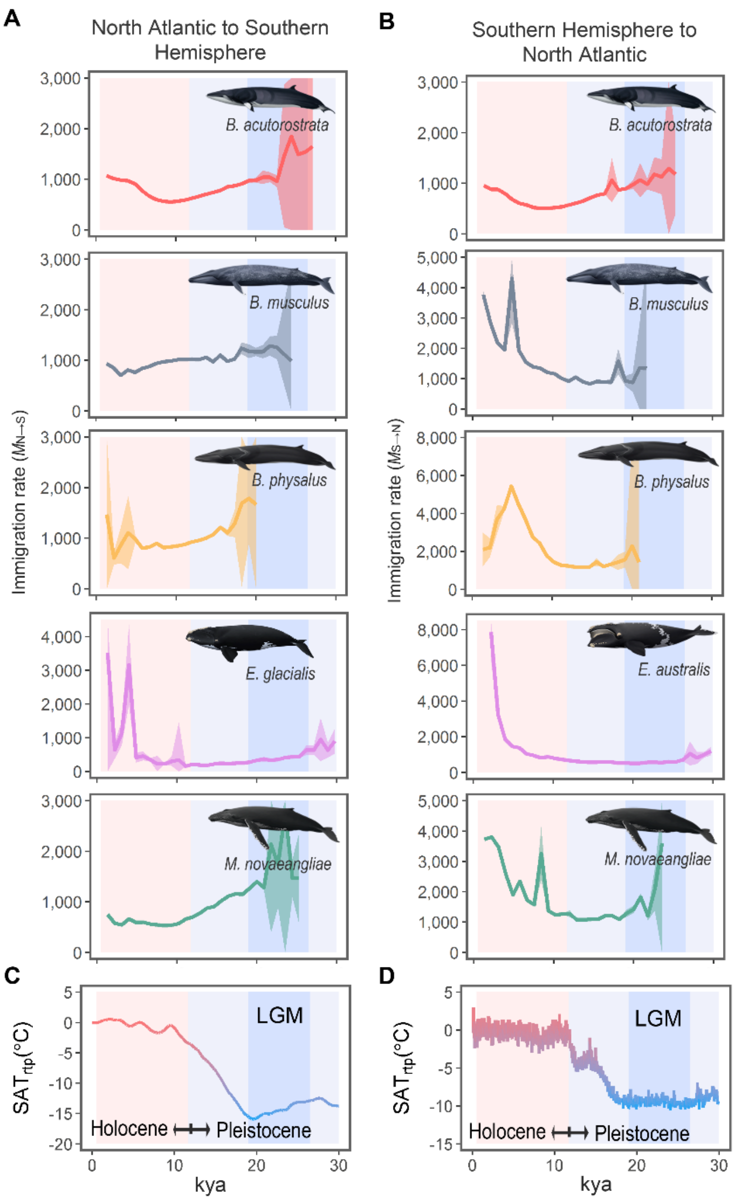


**Fig. S2.**

**Estimates of temporal trends in *M* between North Atlantic Ocean and Southern Hemisphere con-specific baleen whale populations.**

Description: (A) Immigration rate from North to South ($M_{N>S}$) and (B) immigration rate from South to North ($M_{S>N}$). Note different scales for $M$. (C - D) Historical surface air temperature relative to present temperature (SAT$_{RTP}$) in degrees Celsius (°C) for the Northern (C) and Southern Hemisphere (D). Kya: thousands of years ago.

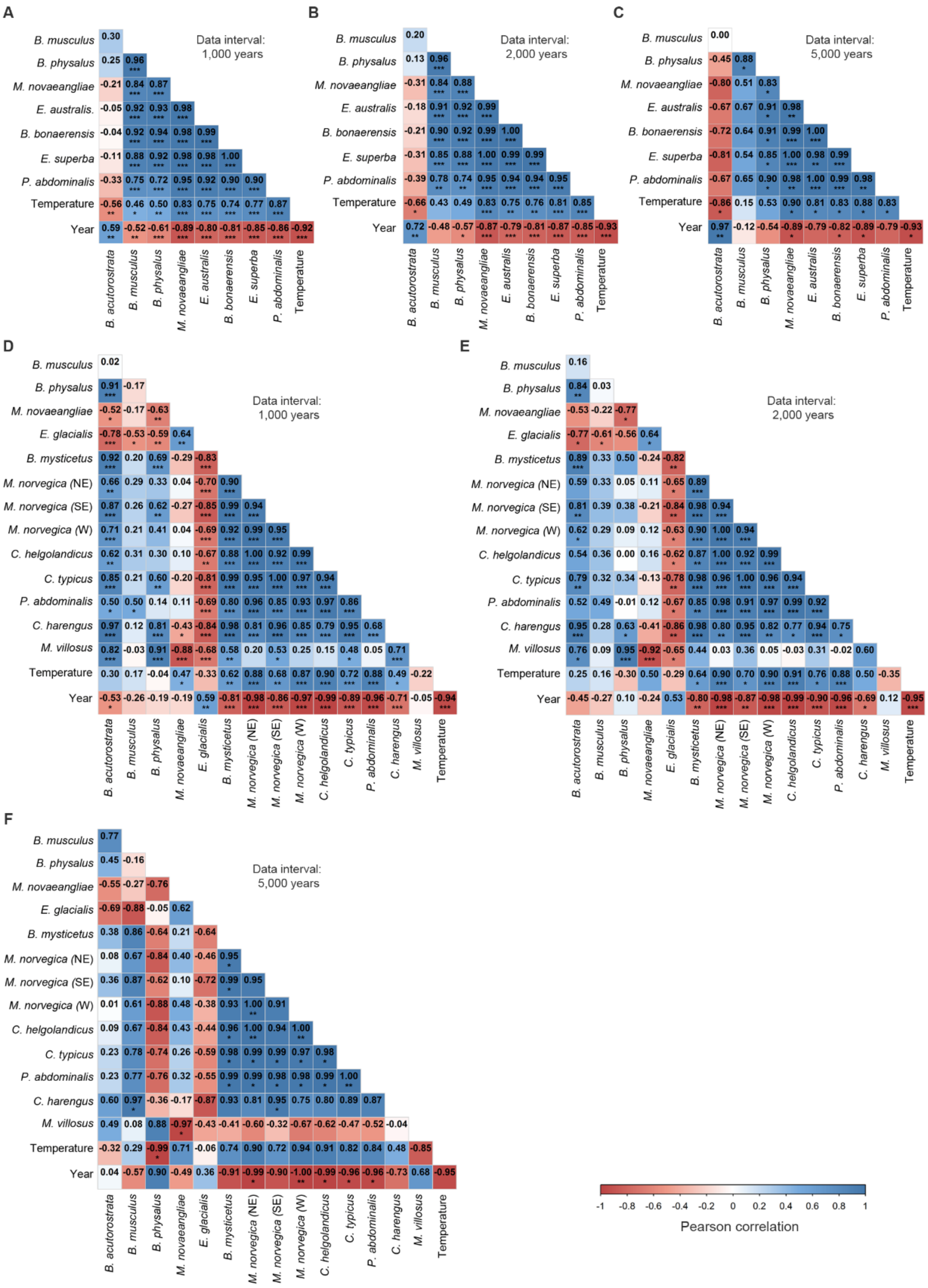


**Fig. S3.**

**Pairwise Pearson's correlations among baleen whale and prey estimates of $\theta$, temperature changes and time.** (A - C) Southern Hemisphere, (D - F) North Atlantic Ocean. Blue: positive correlation, red: negative correlation. *, **, *** denotes p-values below 0.05, 0.005, 0.0005, respectively. Time interval between observations was 1,000, 2,000 or 5,000 years.

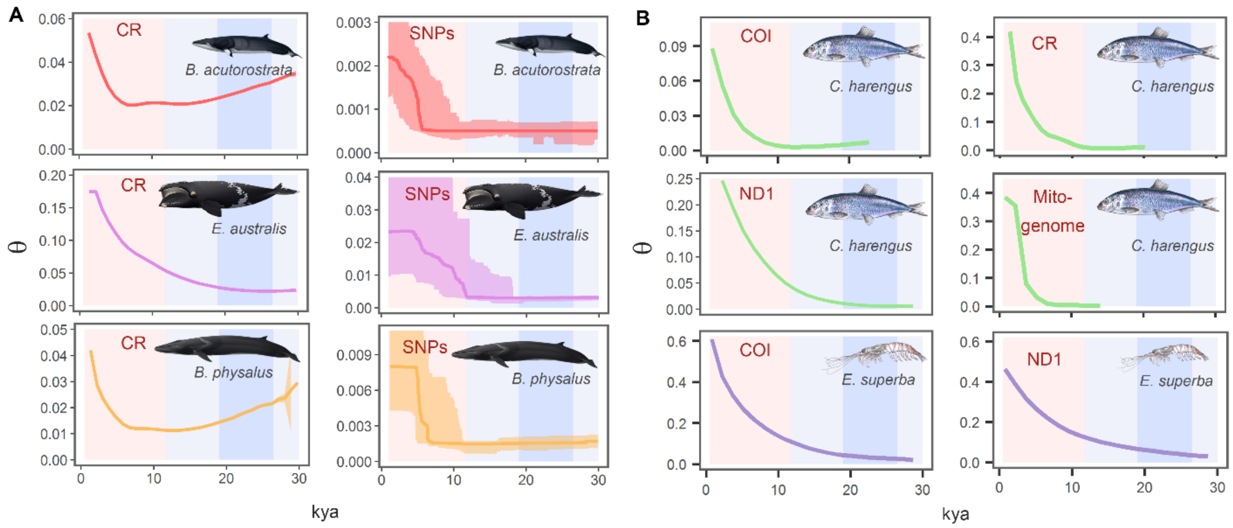


**Fig. S4.**

**Temporal trends in $\theta$ estimated from different mitochondrial genes and genome-wide SNP genotypes.** (A) Temporal change in $\theta$ based upon mitochondrial and nuclear DNA data in common minke whale, southern right whale and fin whale. (B) Estimated change in $\theta$ based upon data from different mitochondrial genes in North Atlantic herring and Antarctic krill. CR: mtDNA control region, mtDNA COI: cytochrome *c* oxidase subunit 1, mtDNA ND1: NADH dehydrogenase subunit 1. Mitogenome: entire mitochondrial genome. SNPs: genome-wide single nucleotide polymorphism genotypes at a minimum coverage of x2. The red- and light blue-shaded areas represent the Holocene and Pleistocene period, respectively. The dark blue-shaded area indicates the LGM. Kya: thousands of years ago. Note the different scales of the vertical axis.

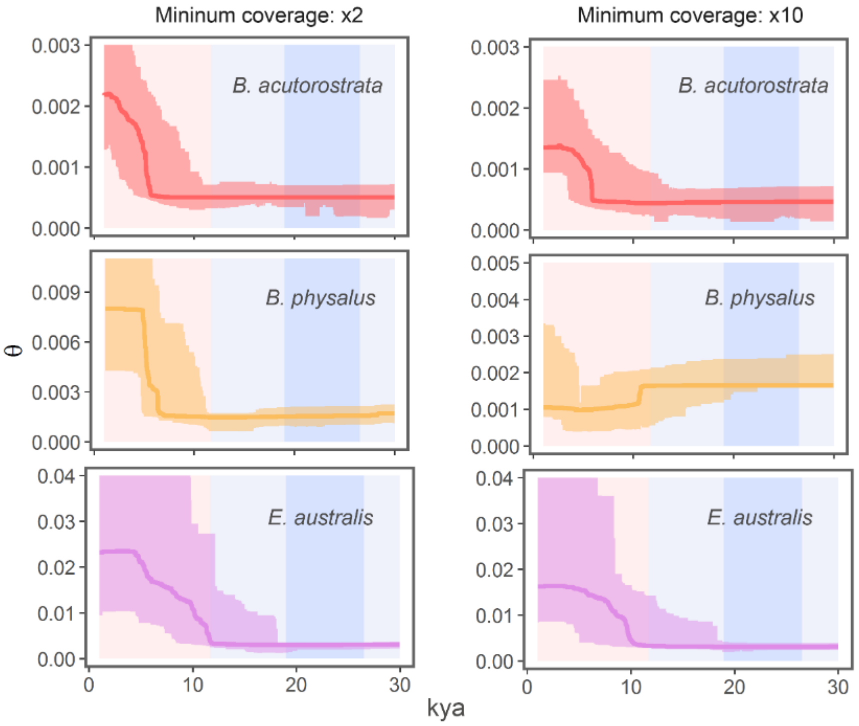


**Fig. S5.**

**Temporal trends in $\theta$ estimated from genome-wide SNP genotypes of two different levels of minimum coverage.**

Estimated demographic history employing genome-wide SNP genotypes with a minimum coverage at x2 and x10 in North Atlantic common minke whale (top), North Atlantic fin whale (middle) and southern right whale (bottom). Values on the X-axis denote the time in thousands of years ago (kya) and the Y-axis the estimate of $\theta$.

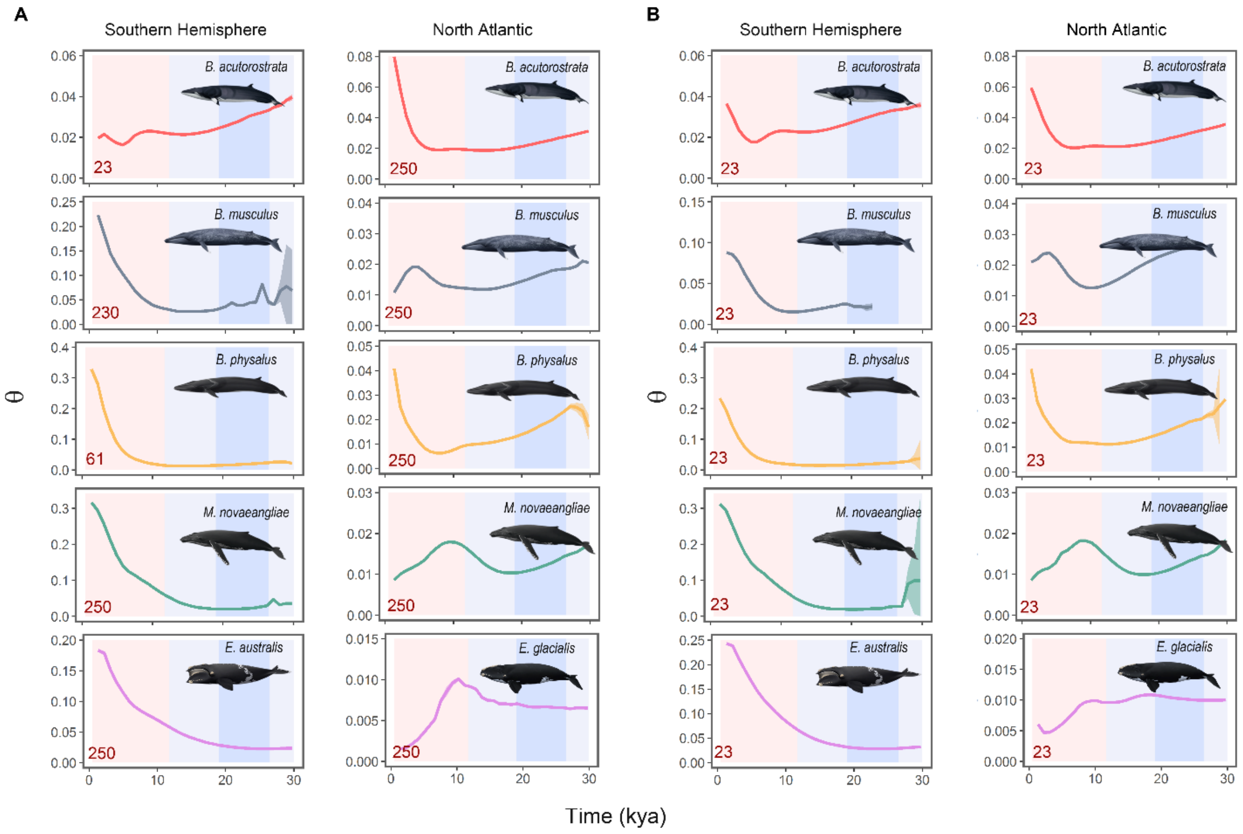


**Fig. S6.**

**Temporal trends in $\theta$ estimated from two different values of random sub-sampling of mitochondrial DNA sequences.**

Temporal change in $\theta$ estimated in baleen whales from a maximum random sub-sampling at: (A) 250 mitochondrial DNA sequences and (B) 23 mitochondrial DNA sequences. Values on the X-axis denote the time in thousands of years ago (kya) and the Y-axis the estimate of $\theta$. The red number at the bottom-right of each plot denotes the sample size employed for each estimation. Note the different scales of the values on the vertical axis in genetic diversity ($\theta$), and the uneven sample size of some populations.

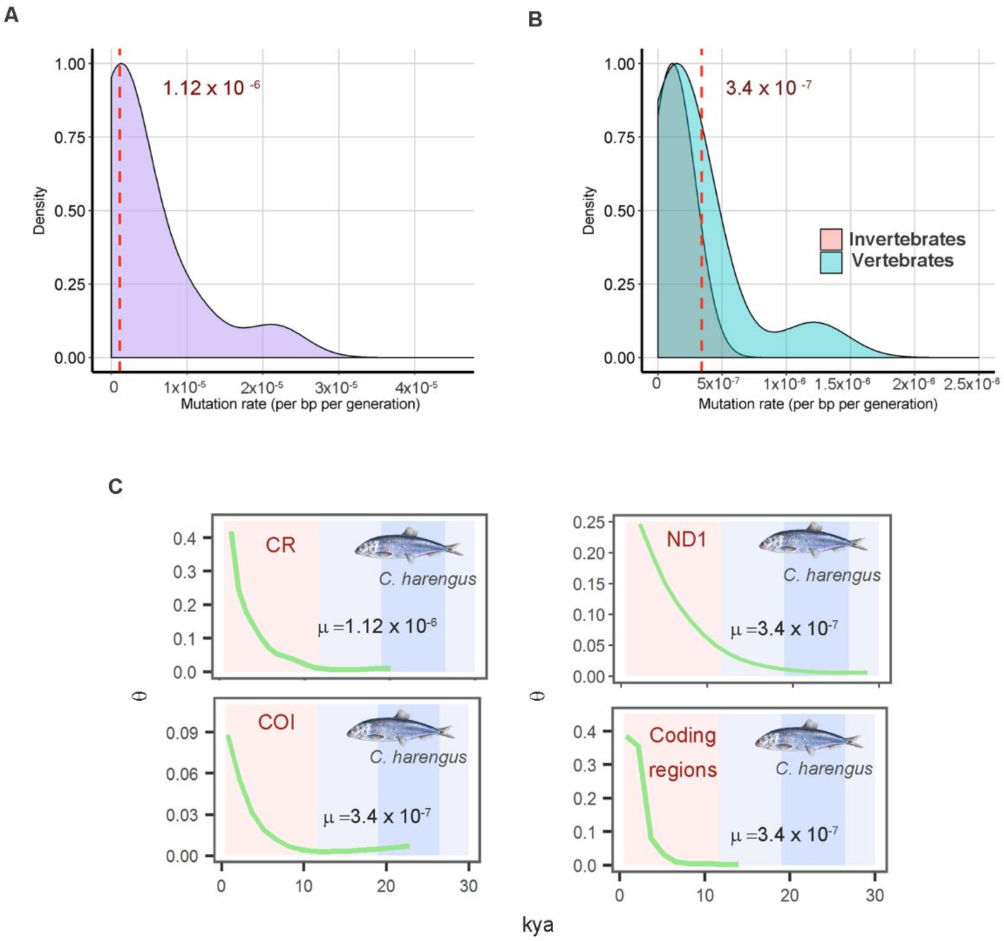


**Fig. S7.**

**Distribution of reported generational mutation rate estimates for coding and non-coding mtDNA and comparison of the trend and timing of change in $\theta$.**

Distribution of reported generational mutation rate estimates for (A) mtDNA control region (non-coding), (B) coding mtDNA (including entire genome) for vertebrates and invertebrates. Distributions based upon the mutation rates listed in Table S1.1-2. The Y-axis denotes the probability of the X-values based on a kernel density estimation. The red vertical line and inserted rate denotes the chosen mutation rates. (C) Estimated demographic history employing different mitochondrial genes in North Atlantic herring (*Clupea harengus*). Values on the horizontal axis denote the time in thousands of years ago (kya). Vertical axis: estimates of $\theta$. Note the different scales of the vertical axis. CR: mtDNA control region, COI: mtDNA cytochrome *c* oxidase, subunit 1, ND1: mtDNA NADH dehydrogenase, subunit 1. Coding regions: entire mitochondrial genome excluding the control region. $\mu$: the generational mutation rate applied in the analysis. The red- and light blue-shaded areas represent the Holocene and Pleistocene period, respectively. The dark blue-shaded area indicates the LGM.

**Table S1.**

**Species list and sample collection for the North Atlantic and Southern Hemisphere**.

| Species | Common name | Sampling region | n | Marker | Sequence length | Source |
|---|---|---|---|---|---|---|
| | | | **North Atlantic** | | | |
| **Baleen whales** | | | | | | |
| *Balaenoptera acutorostrata* | Common minke whale | NA | 931 | CR | 322 | This study |
| *Balaenoptera musculus* | Blue whale | NA | 325 | CR | 404 | This study |
| *Balaenoptera physalus* | Fin whale | WNA | 280 | CR | 391 | This study, (*41*) |
| *Megaptera novaeangliae* | Humpback whale | WI | 1086 | CR | 396 | This study |
| *Eubalaena glacialis* | North Atlantic right whale | WNA | 269 | CR | 381 | (*42*) |
| *Balaena mysticetus* | Bowhead whale | WNA | 395 | CR | 454 | This study |
| **Prey species** | | | | | | |
| *Meganyctiphanes norvegica* | Northern krill | NA | 834* | ND1 | 155 | (*43, 44*) |
| *Calanus helgolandicus* | Copepod | ENA | 218 | 16S | 408 | (*45*) |
| *Centropages typicus* | Copepod | NA | 79 | COI | 560 | (*46*) |
| *Pleuromamma abdominalis* | Copepod | NA | 130 | COI | 441 | (*47*) |
| *Clupea harengus* | Atlantic herring | ENA | 98 | COI | 1,551 | (*48*) |
| *Mallotus villosus* | Capelin | WNA | 41 | CYTB | 572 | (*54*) |
| | | | **Southern Hemisphere** | | | |
| **Baleen whales** | | | | | | |
| *Balaenoptera acutorostrata* | Common minke whale | WSA, SO | 23 | CR | 322 | (*49*) |
| *Balaenoptera musculus* | Blue whale | SO | 230 | CR | 404 | (*51, 112*) |
| *Balaenoptera physalus* | Fin whale | SO | 61 | CR | 391 | This study, (*41*) |
| *Megaptera novaeangliae* | Humpback whale | SA | 500 | CR | 396 | (*33*) |
| *Eubalaena australis* | Southern right whale | SA | 481 | CR | 381 | This study, (*52*) |
| *Balaenoptera bonaerensis* | Antarctic minke whale | WSA, SO | 180 | CR | 337 | (*49*) |
| **Prey species** | | | | | | |
| *Euphausia superba* | Antarctic krill | SO | 640 | COI | 593 | (*53, 113*) |
| *Pleuromamma abdominalis* | Copepod | SA, WIO | 231 | COI | 441 | (*47*) |

Notes: List of species analyzed, sampling region, number of samples (n), molecular marker, sequence length in number of bp, and source. CR: control region, COI: cytochrome *c* oxidase, subunit 1, ND1: NADH dehydrogenase, subunit 1, CYTB: cytochrome *b* and 16S: 16S rDNA of the mtDNA. NA: North Atlantic (NA), ENA: Eastern NA, WNA: Western NA, WI: West Indies, SA: South Atlantic (SA), WSA: Western SA, SO: Southern Ocean, WIO: Western Indian Ocean. *Includes 654 sequences from the Northeastern NA (NE-NA), 146 from the Southeastern NA (SE-NA) and 34 from the Western NA (W-NA).

**Table S2.**

**Prior distributions of estimation parameters in MIGRATE-N.**

| Species | Marker | ti:tv | Parameter $\theta$ | | | | Parameter $M$ | | | |
|---|---|---|---|---|---|---|---|---|---|---|
| | | | Priors | | Starting parameters | | Priors | | Starting parameters | |
| | | | Maximum | Delta | $\theta_S$ | $\theta_N$ | Maximum | Delta | $M_{N>S}$ | $M_{S>N}$ |
| *B. acutorostrata* | CR | 7.7 | 0.1 | 0.01 | 0.01 | 0.03 | 200 | 20 | 50 | 1 |
| *B. musculus* | CR | 19.7 | 0.1 | 0.01 | 0.06 | 0.01 | 300 | 30 | 95 | 1 |
| *B. physalus* | CR | 19.8 | 0.2 | 0.02 | 0.09 | 0.02 | 100 | 10 | 0 | 20 |
| *M. novaeangliae* | CR | 25.4 | 0.15 | 0.015 | 0.07 | 0.007 | 250 | 25 | 60 | 20 |
| *E. glacialis* | CR | 119.5 | 0.1 | 0.01 | 0.04 | 0.003 | 300 | 30 | 1 | 25 |
| *B. mysticetus* | CR | 9.6 | 0.1 | 0.01 | | 0.035 | | | | |
| *B. borealis* | CR | 25.2 | 0.4 | 0.04 | 0.22 | | | | | |
| *E. superba* | COI | 8.2 | 0.5 | 0.05 | 0.19 | | | | | |
| *M. norvegica* | ND1 | 5 | 0.1 | 0.01 | | 0.02 (NE-NA)<br>0.017 (SE-NA)<br>0.012 (W-NA) | | | | |
| *C. helgolandicus* | 16S | 6.5 | 0.1 | 0.01 | | 0.02 | | | | |
| *C. typicus* | COI | 39.2 | 0.5 | 0.05 | | 0.2 | | | | |
| *P. abdominalis* | COI | 6.8 | 0.2 | 0.02 | 0.12 | 0.12 | 500 | 50 | 83 | 83 |
| *C. harengus* | COI | 12.3 | 0.2 | 0.02 | | 0.036 | | | | |
| *M. villosus* | CYTB | 7.4 | 0.05 | 0.005 | | 0.02 | | | | |
| *C. harengus* | CR | 5.6 | 0.4 | 0.04 | | 0.16 | | | | |
| *C. harengus* | ND1 | 12.3 | 0.2 | 0.02 | | 0.04 | | | | |
| *C. harengus* | Mitogenome | 11.3 | 0.2 | 0.02 | | 0.08 | | | | |
| *E. superba* | ND1 | 7.7 | 0.3 | 0.03 | 0.15 | | | | | |

Notes: CR: control region, COI: cytochrome *c* oxidase, subunit 1, ND1: NADH dehydrogenase, subunit 1, CYTB: cytochrome *b*, 16S: 16S rDNA of the mtDNA and mitogenome: entire mitochondrial genome excluding the control region. The transition:transversion rate (ti:tv). The prior parameters and the starting parameter values are shown for $\theta$ and $M$. A uniform distribution and a minimum prior of zero were employed for all priors. $\theta_N$: $\theta$ North Atlantic population, $\theta_S$: $\theta$ Southern Hemisphere population, $M_{N>S}$: immigration rate from the North Atlantic Ocean into the Southern Hemisphere and $M_{S>N}$: immigration rate from the Southern Hemisphere into North Atlantic Ocean. See Table S1 for the definition of the abbreviations: NE-NA, SE-NA and W-NA.

**Table S3.**
**Species and sample sizes.**

| Species | Common name | Sampling region | Marker | n | Sequence length/Number of SNPs | Source |
|---|---|---|---|---|---|---|
| *C. harengus* | Atlantic herring | ENA | COI | 98 | 1,551 | (*48*) |
| *C. harengus* | Atlantic herring | ENA | CR | 98 | 1,055 | (*48*) |
| *C. harengus* | Atlantic herring | ENA | ND1 | 98 | 975 | (*48*) |
| *C. harengus* | Atlantic herring | ENA | Mitogenome | 98 | 15,653 | (*48*) |
| *E. superba* | Antarctic krill | SO | COI | 640 | 593 | (*53, 113*) |
| *E. superba* | Antarctic krill | SO | NDI | 139 | 494 | (*113*) |
| *B. acutorostrata* | Common minke whale | NA | CR | 867 | 322 | This study |
| *B. acutorostrata* | Common minke whale | NA | SNPs | 27 | 14,304 (x10)<br>24,988 (x2) | This study |
| *E. australis* | Southern right whale | SA | CR | 481 | 381 | This study, (*52*) |
| *E. australis* | Southern right whale | ESA | SNPs | 45 | 31,482 (x10)<br>68,575 (x2) | This study |
| *B. physalus* | Fin whale | WNA | CR | 280 | 391 | This study |
| *B. physalus* | Fin whale | NA | SNPs | 28 | 29,544 (x10)<br>56,325 (x2) | This study |

Notes: Species, sampling region, molecular marker, sample size (n), sequence length in number of bp or number of estimated SNPs (i.e., number of inferred polymorphic sites from the site frequency spectrum) for each species with minimum coverage at x10 and x2. CR: control region, COI: the cytochrome *c* oxidase subunit 1, ND1: the NADH dehydrogenase subunit 1 of the mtDNA, Mitogenome: entire mitochondrial genome excluding the control region, SNPs: single nucleotide polymorphism genotypes. See Table S1 for the definition of the abbreviations: NA, ENA, WNA, SA, SO and ESA.

**Table S4.**

**Summary of mutation rates estimated for the mitochondrial DNA control region sequence.**

| Marker | Species | Mutation rate bp/year (95 % CI) | Reference | Generation time | Mutation rate bp/generation (95 % CI) |
|---|---|---|---|---|---|
| CR | *Eschrichtius robustus* | $5.40x10^{-8}$ ($4.96x10^{-8}$ - $6.16x10^{-8}$) | (*65*) | 25.76 | **$1.39x10^{-6}$** ($1.28x10^{-6}$ - $1.59x10^{-6}$) |
| CR | *Eschrichtius robustus* | 4.80x10-$^{8}$ ($4.32x10^{-8}$ - $5.36x10^{-8}$) | (*65*) | 25.76 | **$1.24x10^{-6}$** ($1.11x10^{-6}$ - $1.38x10^{-6}$) |
| CR | *Balaenoptera acutorostrata* | $5.00x10^{-08}$ ($4.70x10^{-8}$ - $5.39x10^{-8}$) | (*65*) | 21.22 | **$1.06x10^{-6}$** ($9.97x10^{-7}$ - $1.14x10^{-6}$) |
| CR | *Balaenoptera acutorostrata* | $5.30x10^{-8}$ ($4.94x10^{-8}$ - $5.66x10^{-8}$) | (*65*) | 21.22 | **$1.12x10^{-6}$** ($1.05x10^{-6}$ - $1.20x10^{-6}$) |
| CR | *Balaenoptera acutorostrata* | $4.40x10^{-8}$ ($4.00x10^{-8}$ - $4.69x10^{-8}$) | (*65*) | 21.22 | **$9.34x10^{-7}$** ($8.49x10^{-7}$ - $9.95x10^{-7}$) |
| CR | *Megaptera novaeangliae* | $4.60x10^{-8}$ ($3.59x10^{-8}$ - $5.50x10^{-8}$) | (*65*) | 26.87 | **$1.24x10^{-6}$** ($9.65x10^{-7}$ - $1.48x10^{-6}$) |
| CR | *Megaptera novaeangliae* | $5.20x10^{-8}$ ($4.12x10^{-8}$ - $6.32x10^{-8}$) | (*65*) | 26.87 | **$1.40x10^{-6}$** ($1.11x10^{-6}$ - $1.70x10^{-6}$) |
| CR | *Megaptera novaeangliae* | $8.50x10^{-9}$ ($7.0x10^{-9}$ – $1.0x10^{-8}$) | (*114*) | 26.87 | **$2.28x10^{-7}$** ($1.88x10^{-7}$ - $2.69x10^{-7}$) |
| CR | *Balaena mysticetus* | $1.50x10^{-7}$ (n/a) | (*76*) | 52.67 | **$7.90x10^{-6}$** (n/a) |
| CR | *Balaena mysticetus* | $2.10x10^{-7}$ ($1.22x10^{-7}$ - $3.03x10^{-7}$) | (*76*) | 52.67 | **$1.11x10^{-5}$** ($6.43x10^{-6}$ - $1.60x10^{-5}$) |
| CR | *Balaena mysticetus* | $2.00x10^{-8}$ ($1.20x10^{-8}$ - $3.70x10^{-8}$) | (*77*) | 52.67 | **$1.05x10^{-6}$** ($6.32x10^{-7}$ - $1.95x10^{-6}$) |
| CR | *Balaena mysticetus* | $4.11x10^{-7}$ ($2.07x10^{-7}$ - $6.49x10^{-7}$) | (*115*) | 52.67 | **$2.16x10^{-5}$** ($1.09x10^{-5}$ - $3.42x10^{-5}$) |

Notes: Mutation rates per base pair per year (bp/year) and per generation (bp/generation). CR: mtDNA control region. Numbers in parentheses denote the 95% confidence interval of the mutation rate estimates. Generation time represents the average estimates reported by Pacifici *et al*. (*66*) and Taylor *et al.* (*67*). n/a: not available.

**Table S5.**

**Summary of mutation rates estimated for the coding genes of the mitochondrial DNA and for the entire mitochondrial genome.**

| Marker | Species | Mutation rate bp/year (95 % CI) | Reference | Generation time | Mutation rate bp/generation (95 % CI) | Reference |
|---|---|---|---|---|---|---|
| **Vertebrates** | | | | | | |
| CYTB | *Eschrichtius robustus* | $4.00x10^{-9}$ ($3.87x10^{-9}$ - $4.13x10^{-9}$) | (*65*) | 25.76 (*66, 67*) | **$1.03x10^{-7}$** ($9.97x10^{-8}$ - $1.06x10^{-7}$) | |
| CYTB | *Balaenoptera borealis* | $7.00x10^{-9}$ ($3.00x10^{-9}$ - $1.20x10^{-8}$) | (*116*) | 22.05 (*66, 67*) | **$1.54x10^{-7}$** ($6.62x10^{-8}$ - $2.65x10^{-7}$) | |
| CYTB | *Megaptera novaeangliae* | $8.00x10^{-9}$ ($5.00x10^{-9}$ - $2.00x10^{-8}$) | (*116*) | 26.87 (*66, 67*) | **$2.15x10^{-7}$** ($1.34x10^{-7}$ - $5.37x10^{-7}$) | |
| Mitogenome | *Ursus maritimus* | $1.12x10^{-8}$ ($8.96x10^{-9}$ - $1.50x10^{-8}$) | (*117*) | 15 (*66*) | **$1.68x10^{-7}$** ($1.34x10^{-7}$ - $2.25x10^{-7}$) | |
| Mitogenome | *Orcinus orca* | $2.60x10^{-9}$ ($1.50x10^{-9}$ - $3.83x10^{-9}$) | (*118*) | 31.8 (*66, 67*) | **$8.27x10^{-8}$** ($4.77x10^{-8}$ - $1.22x10^{-7}$) | |
| Coding mtDNA | *Homo sapiens* | $1.70x10^{-8}$ (n/a) | (*68*) | 20 (*69*) | **$3.40x10^{-7}$** (n/a) | |
| Coding mtDNA | *Homo sapiens* | $1.26x10^{-8}$ ($1.18x10^{-08}$ - $1.34x10^{-8}$) | (*119*) | 20 (*69*) | **$2.52x10^{-7}$** ($2.36x10^{-7}$ - $2.68x10^{-7}$) | |
| Coding mtDNA | *Homo sapiens* | $6.09 x10^{-8}$ (n/a) | (*78*) | 20 (*69*) | **$1.22 x10^{-6}$** (n/a) | |
| CYTB | *Anisotremus* spp. | | | | **$1.60x10^{-8}$** ($1.50x10^{-8}$ - $1.70x10^{-8}$) | (*120*) |
| Mitogenome | *Clupea harengus* | $4.70x10^{-9}$ ($3.50x10^{-9}$ - $6.00x10^{-9}$) | (*48*) | 6.5 (*121*) | **$3.06x10^{-8}$** ($2.28x10^{-8}$ - $3.90x10^{-8}$) | |
| **Invertebrates** | | | | | | |
| COI | *Melarhaphe neritoides* | $5.82x10^{-5}$ | (*122*) | 3.42 (*122*) | **$1.99x10^{-4}$** | (*122*) |
| COI | *Mytilus edulis* | | | | **$9.51x10^{-8}$** ($5.67x10^{-8}$ – $1.34x10^{-7}$) | (*123*) |
| COI | *Asteria rubens* | | | | **$4.84 x10^{-8}$** ($3.12x10^{-8}$ – $6.56x10^{-8}$) | (*123*) |
| COI | *Nucella lapillus* | | | | **$4.43 x10^{-8}$** ($2.33x10^{-8}$ – $6.53x10^{-8}$) | (*123*) |
| COI | *Littorina obtusata* | | | | **$2.49 x10^{-8}$** (0.00 – $5.04x10^{-8}$) | (*123*) |
| COI | *Semibalanus balanoides* | | | | **$2.76 x10^{-8}$** ($1.52x10^{-8}$ – $4.00x10^{-8}$) | (*123*) |
| CYTB | *Paralithodes camtschaticus* | $5.00x10^{-9}$ (n/a) | (*124*) | 5 (*124*) | **$2.50x10^{-8}$** (n/a) | |
| Mitogenome | *Daphnia pulex* | | | | **$1.73x10^{-7}$** (n/a) | (*71*) |
| Mitogenome | *Daphnia pulex* | | | | **$1.37x10^{-7}$** (n/a) | (*71*) |
| Mitogenome | *Caenorhabditis elegans* | | | | **$1.60x10^{-7}$** ($1.29x10^{-7}$ - $1.91x10^{-7}$) | (*70*) |
| Mitogenome | *Drosophila melanogaster* | | | | **$6.20x10^{-8}$** ($3.00x10^{-8}$ - 1.14x10-$^{7}$) | (*125*) |

Notes: Estimates are in mutations per base pair per year (bp/year) and mutations per generation (bp/generation). COI: cytochrome c oxidase, subunit I, CYTB: cytochrome *b* of the mtDNA, Coding mtDNA: all coding DNA sequences in the mtDNA genome (i.e., excluding non-coding region). Mitogenome: entire mtDNA genome. Numbers in parentheses denote the 95% confidence interval of the estimated mutation rate. Reference of the mutation rate per bp/year or bp/generation. Generation time: average generation length estimate. n/a: not available.
